# Unravelling the complex biogeographic and anthropogenic history of Alaska’s mountain goats

**DOI:** 10.1101/2023.08.07.552341

**Authors:** Kiana B. Young, Kevin S. White, Aaron B.A. Shafer

## Abstract

**Aim:** We used genetic tools to examine the population structure of mountain goats in Alaska, USA and assessed the demographic history of this species in relation to the natural and anthropogenic forces.

**Location:** Alaska, USA

**Taxon:** North American mountain goat (*Oreamnos americanus*)

**Methods:** Samples were collected between 2006 - 2020 from harvested animals and live captures. We genotyped 816 mountain goats at 18 microsatellite loci and identified the number of genetically distinct subpopulations across the state and assessed their genetic diversity. We used Bayesian computation software to investigate the demographic history relative to the known biogeographic history of the state. We also simulated island translocation events and compared simulations to empirical data to address the hypothesis that Baranof Island was a cryptic refugia.

**Results:** We showed that Alaska has four genetically distinct subpopulations of mountain goats with some additional genetic structure within those subpopulations. The main split of mountain goats between Southcentral and Southeast Alaska occurred ∼14,000 years ago. Simulations of translocation events largely aligned with the current populations observed today except for Baranof Island which showed greater diversity than the translocation simulation.

**Main Conclusions:** The distribution and genetic structure of mountain goats in Alaska reflects a combination of natural and anthropogenic forces. A rapid northerly expansion through an ice-free corridor in combination with the isolated nature of the landscape led to low diversity and isolation 14,000 years ago in Southcentral Alaska and higher diversity in Southeast Alaska. Two of the three islands where mountain goat translocations have occurred match genetically with their source population, while Baranof Island appears to have a divergent population, consistent with the hypothesis of an endemic or cryptic population prior to the translocation event. This study highlights the value of considering both the natural and anthropogenic forces when assessing the biogeographic history of a species.

## Introduction

Alaska has a dynamic geologic and anthropogenic history that has impacted contemporary wildlife populations. The state is geographically diverse, discontinuous from the majority of the United States – colloquially referred to as the “Lower 48” – and is characterized by rugged mountainous terrain, deep glacier- and river-formed valleys, long complex coastlines and archipelagos, and numerous glaciers and icefields (Brooks et al., 1906; Stowell, 2006). During the Wisconsin glaciation of the Pleistocene Ice Age, the southern portion of Alaska was covered by the Cordilleran Ice Sheet (Clark et al., 2009). Two large scale refugia, Beringia and Pacific Northwest, are widely believed to have been the source of Alaska’s flora and fauna, but evidence for additional smaller-scale coastal refugia exists (Sawyer et al., 2019; Shafer et al., 2010; Shafer, Côté, et al., 2011). Since the Wisconsin glaciation, smaller scale ice age events have occurred in Alaska such as the Little Ice Age (270 years ago) in Southeast Alaska (Connor et al., 2009). While these events did not have the same magnitude of impact on biodiversity as the large-scale events (Mann & Streveler, 2008), they are still relevant when examining the composition of plants and animals across the landscape.

Beyond the naturally occurring geologic events, humans have also had an impact on Alaska’s biodiversity. Indigenous peoples have inhabited the landscape for thousands of years, relying on plants and animals for food, clothing, tools, and a variety of other uses (Dixon, 2001). More recently, the European colonization and subsequent development in Alaska has expanded the human impacts on these wildlife populations. Human development has grown and expanded and the demand for harvest opportunities has increased which led to a period of translocation events of certain high-value species to areas where they did not exist prior (Paul, 2009). The combination of both natural and anthropogenic impacts on wildlife is key for interpreting the current structure of a population and offering insight necessary for ensuring the future of a species that is valued for both its place in the ecosystem and its significance to human cultures.

Mountain goats (*Oreamnos americanus*) principally inhabit coastal mountain ranges throughout the southeastern and southcentral portions of Alaska (Côté & Festa-Bianchet, 2008; MacDonald & Cook, 2009). Mountain goats are habitat specialists and utilize steep, rugged terrain - an adaptation to risk of predation from large carnivores (Côté & Festa-Bianchet, 2003; Fox & Streveler, 1986). They exhibit morphological adaptations to their mountainous habitat including hooves and body shape that enable navigation of steep, rocky terrain and thick, white fur that facilitates thermoregulation in extreme winter climates and camouflages them in their high elevation, often snowy habitat (Côté & Festa-Bianchet, 2008; Déry et al., 2019). Mountain goats are among the most culturally important species in Alaska for both hunting and wildlife viewing, and are of special interest from a biogeographical and anthropogenic perspective because of their unique habitat associations (Shafer et al., 2012), indigenous cultural traditions (notably Chilkat and Raven’s tail robes; Rofkar, 2014), and management interventions (i.e. translocations and harvest; Mcdonough and Selinger, 2008; Paul, 2009; White, Levi, et al., 2021).

Mountain goats originally colonized Beringia from Asia and, during the various ice ages, were isolated into northern, southern, and coastal refugial populations (Martchenko & Shafer, 2022; Nagorsen & Keddie, 2000; Shafer, White, et al., 2011). Following the ice age and recession of glacial barriers, populations expanded to their present distribution (Shafer, Côté, et al., 2011). However, population translocations occurred over the last 100 years (Paul, 2009) and altered distribution and presumably population genetic structure (Shafer, White, et al. 2011). Mountain goats were successfully translocated to three islands in Alaska: Baranof Island, Kodiak Island, and Revillagigedo (Revilla) Island. A few other occasions of mountain goat translocations occurred in the state, but they were either unsuccessful or considered augmentations rather than full translocations. Today, each of these successful introduced groups of mountain goats have established themselves as productive populations (Paul, 2009), but little is known about the genetic effects of these translocations. Genetic drift and inbreeding have a greater impact on populations when their size and amount of gene flow are reduced (Frankham, 1995, 1996), as is the case with islands. Of the three island populations, Baranof Island mountain goats are of special interest because previous genetic research has suggested that an endemic or cryptic population of mountain goats could have existed on Baranof Island before individuals were introduced (Shafer, White, et al., 2011).

Here, we used an extensive genotype dataset and simulations to investigate the biogeographical history and current population structure of mountain goats in Alaska. Our objectives for this study were to: 1) infer the demographic history of Alaska mountain goats relative to the known geologic history of the region; 2) identify distinct subpopulations of mountain goats across the state and assess the genetic diversity of each subpopulation; and 3) simulate island translocation events to evaluate the possibility of a Baranof Island cryptic refugia.

## Methods

### Study area and sampling

There are an estimated 24,000-33,500 mountain goats in Alaska, found on the eastern panhandle, Baranof Island, and along the coast of Southcentral Alaska to Kodiak Island (Côté & Festa-Bianchet, 2008). We collected samples between 2006 and 2020 from hunter-harvested animals and live-capture studies. For hunter harvest samples, tissue was collected by Alaska Department of Fish and Game (ADFG) biologists during inspections of hunter harvested animals. In addition, tissue samples were collected from immobilized animals during capture and handling events associated with field research activities (White, Watts, et al., 2021; U.S. Fish & Wildlife Service and ADFG Institutional Animal Care and Use Committees protocols: 2017-006, 05-11, 2016-25, 0078-2018-68, 0039-2017-39). Tissue samples were stored in ethanol, Longmire’s solution, or salt until processed. For each sample, the general location, latitude, longitude, and sex of animal were recorded.

### DNA isolation and genotyping

Tissue samples were digested at 56 °C, and DNA was extracted using a Qiagen DNEasy Blood and Tissue Kit (Qiagen Inc., Valencia, CA, USA) following established protocols (Shafer et al., 2010; Young et al., 2022). We amplified 18 polymorphic microsatellites over three multiplex PCR pools using previously published PCR parameters (White, Levi, et al., 2021).

Samples were genotyped using an ABI 3730 DNA Analyzer (Applied Biosystems, Foster City, California, USA). Positive and negative controls were included to track contamination during extraction and PCR processes. We manually detected and scored loci using the program Geneious (v10.1.2; Kearse et al., 2012). Loci with poor quality DNA (peaks <250 RFUs) were removed, and samples with genotypes less than 80% complete were dropped from the analysis.

### Population structure and diversity

To test for deviations from Hardy-Weinberg Equilibrium (HWE) and linkage disequilibrium (LD), we used the package GENEPOP (version 4.6; Rousset, 2008). Loci that were out of HWE or LD were removed from further analysis. We visualized clusters of samples using a principal component analysis (PCA) and compared principal components to latitude and longitude using linear regression in R (v4.0.4). The Bayesian clustering program STRUCTURE (v2.3.4; Pritchard et al., 2000; Falush et al., 2003, 2007; Hubisz et al., 2009) was used to identify subpopulations within the total population, defined as mountain goats in Alaska. We ran STRUCTURE with independent runs from *K* = 1 to *K* = 25 with 1×10^6^ MCMC, a burn-in period of 1×10^5^, repeated over 20 runs and under correlated and uncorrelated allele frequency scenarios. The most likely number of subpopulations was determined by comparing Puechmaille (2016) and Evanno et al. (2005) methods which use MedMeaK/MaxMeaK/MedMedK/MaxMedK and DeltaK estimators, respectively, and the geographic and known history of the area. Subpopulations were named based on their general location: Southcentral = SOC, northern Southeast = NSE, Baranof Island = BAR, and southern Southeast = SSE. All subsequent analysis used these STRUCTURE subpopulations.

We used the program SPAGeDi (v1.5; Hardy and Vekemans, 2002) to calculate Nei’s *D* and *F_ST_* which we used to assess the genetic differentiation between each of the STRUCTURE defined subpopulations. Isolation-by-distance was calculated in R using the Moran’s I parameter produced by SPAGeDi. We estimated subpopulation-level expected and observed heterozygosity using the package ‘adegenet’ (Jombart, 2008). *F_IS_* was calculated using Genalex (v6.503, (Peakall & Smouse, 2006). We used the program NeEstimator (v2.1; Do et al., 2014) to calculate *N_e_* of each of the subpopulations due to its ability to apply multiple methods of estimating *N_e_* on the same dataset, allowing for a more accurate estimation.

### Demographic history

Based on the subpopulation groupings identified using STRUCTURE, we estimated the timing and patterns of subpopulation divergence events using the approximate Bayesian computational software DIYABC (v2.1.0; Cornuet et al. 2014). DIYABC simulates a dataset and summary statistics for a user-defined demographic scenario and then compares the simulated datasets to the observed dataset. After identifying clusters using the PCA and STRUCTURE analysis, we subsampled 100 individuals from each cluster and ran the demographic analysis. We modeled the scenario of BAR, NNE, and SSE splitting at T1 and SOC splitting at T2 with the objective of determining T2. We simulated 4 x 10^6^ datasets and used the summary statistics: mean number of alleles, genic diversity, and size variance for one- and two-sample summary statistics, mean Garza-Williamson’s M for one sample summary statistic, and *F_ST_* and Classification index, shared allele distance, and (dµ)^2^ distance for two sample summary statistics. We adjusted parameters based on their posterior distribution to further increase the accuracy of the scenario. Using the results from DIYABC, we estimated split time of the original major split of mountain goats in Alaska. Time estimates were reported in number of generations, and we multiplied by a generation time of 6 years following Martchenko et al. (2020).

### Translocation simulations

In 1923, 18 individuals were captured around Tracy Arm in Southeast, Alaska, and transported to Baranof Island, where it was believed that mountain goats did not exist (Paul, 2009). In 1953, 17 mountain goats from the Kenai Peninsula around Seward were released on to Kodiak Island off the coast of Southcentral Alaska. A third translocation event occurred during 1983 and 1991, where a total of 33 individuals were captured around the Quartz Hills area and the Upper Cleveland Peninsula of Southeast Alaska and released onto Revilla Island (Table S1; Paul, 2009).

We simulated these three island translocation events using the software BottleSim (v2.6; Kuo & Janzen, 2003). BottleSim uses an overlapping generation model to simulate the changes in genetic diversity over a given number of years following a population bottleneck. Because we know the exact number of individuals that were translocated, the year that the translocation occurred, and the location from which individuals were taken, we were able to adjust the parameters to match closely with what is observed on those island populations today. For each island, we used genotypes from the source population to represent the translocated individuals (Figure 1). If the sex ratio was known, we used it for the number of females; if the sex ratio was not known, we assumed a ratio of 1:1 for males:females. We used historic survey data and population estimates published by ADFG, when available, to calibrate population growth in the simulations (Table S2). We ran the *Diploid Multilocus with Variable Population Size* model. In the final version of the runs, we set the age of reproduction to 4 and the longevity to 11. For reproductive system, we used the Dioecy with single reproducing male each year. We set the overlapping generations option to 100 which dictates that each individual in generation 0 is randomly assigned an age rather than given an age of zero. Each simulation was run 1000 times, and diversity statistics were averaged over all runs. We calculated the *H_O_*, *F_IS_*, and allelic richness for both the simulated and the observed populations. Allelic richness was calculated using the program ADZE (v1.0; Szpiech et al., 2008). A one sample t-test was conducted to identify the differences in means between the observed and simulated populations.

**Figure 1.**
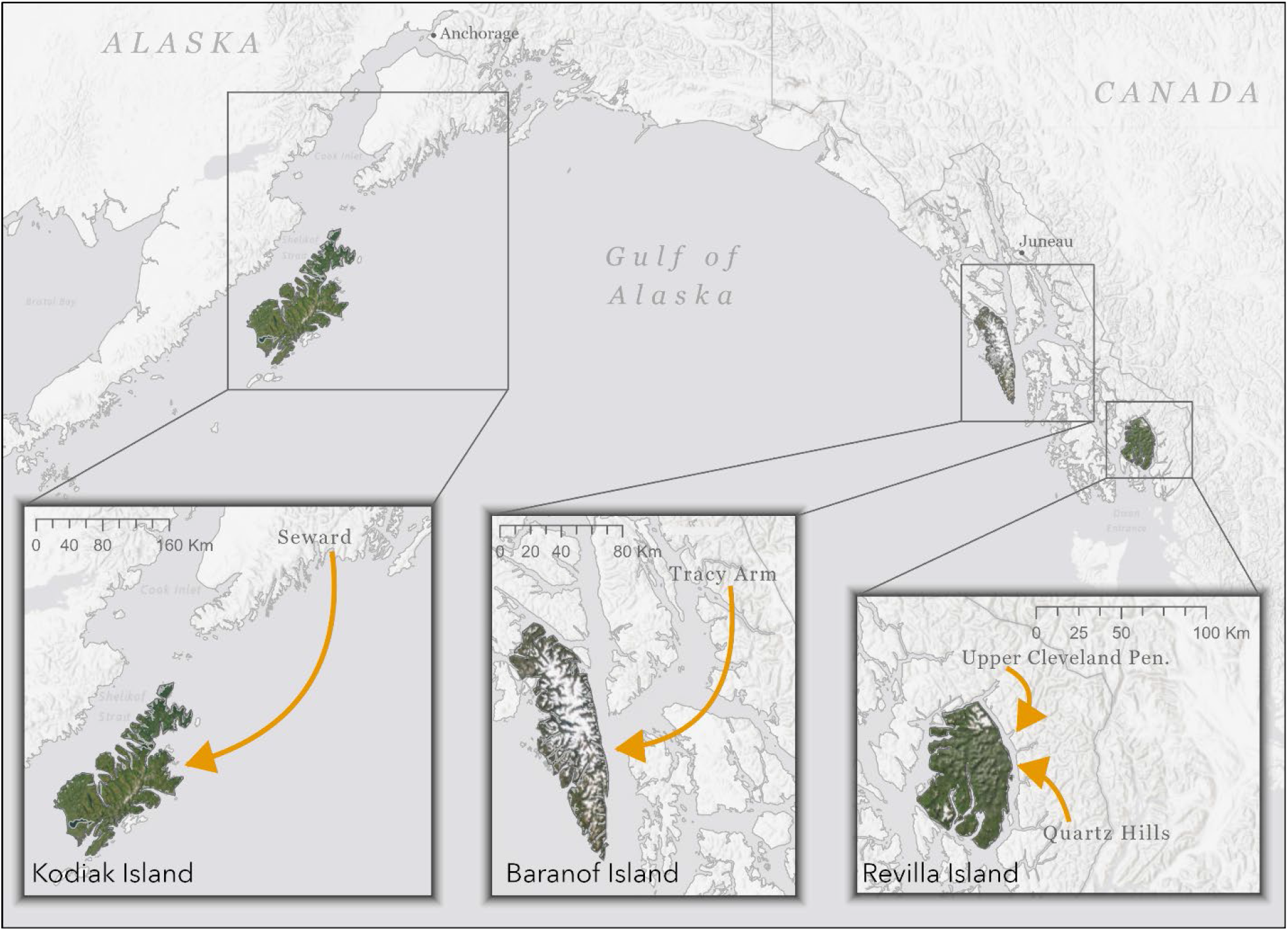
Map of the three island translocation events of mountain goats in Alaska. The source location or locations of the translocated individuals is specified by the arrows.

## Results

### Microsatellite genotyping and population structure

A total of 816 mountain goat samples from across Alaska were genotyped over 18 loci. Genotype completeness for samples ranged from 83%-100% with an average of 98%. PCAs showed broad-scale clustering across all samples in Alaska. Two clear clusters emerged from the PCA, splitting mountain goats into Southcentral and Southeast groupings (Figure 2). STRUCTURE analysis showed additional evidence for a Southcentral and Southeast split in addition to further population structure. The Evanno et al. (2005) Δ*K* estimator indicated that *K* = 2 on a broad scale with finer scale structure present in further hierarchical analysis. The Puechmaille, (2016) method indicated that *K* = 5 (Figure S1). Accordingly, we examined the STRUCTURE assignments and q-values in addition to the most biologically likely groupings based on topography and determined that *K* = 4 in Alaska (Figure 3). Subpopulations are referred to as SOC, NSE, BAR, and SSE based on their location. Hierarchical STRUCTURE analysis was done on each of the four subpopulations, and two of those subpopulations, NSE and SSE, showed additional structure (Figure S2), though the structure was not geographically clustered and distinct enough for us to consider them as fully genetically distinct subpopulations. PC1 vs longitude analysis indicated that clustering existed across a longitudinal gradient (Figure 4; R^2^ = 0.85, p < 0.01). PC1 vs latitude showed a weaker correlation (R^2^ = 0.39, p < 0.01) and PC2 vs latitude showed very little correlation (R^2^ = 0.02, p < 0.01).

**Figure 2.**
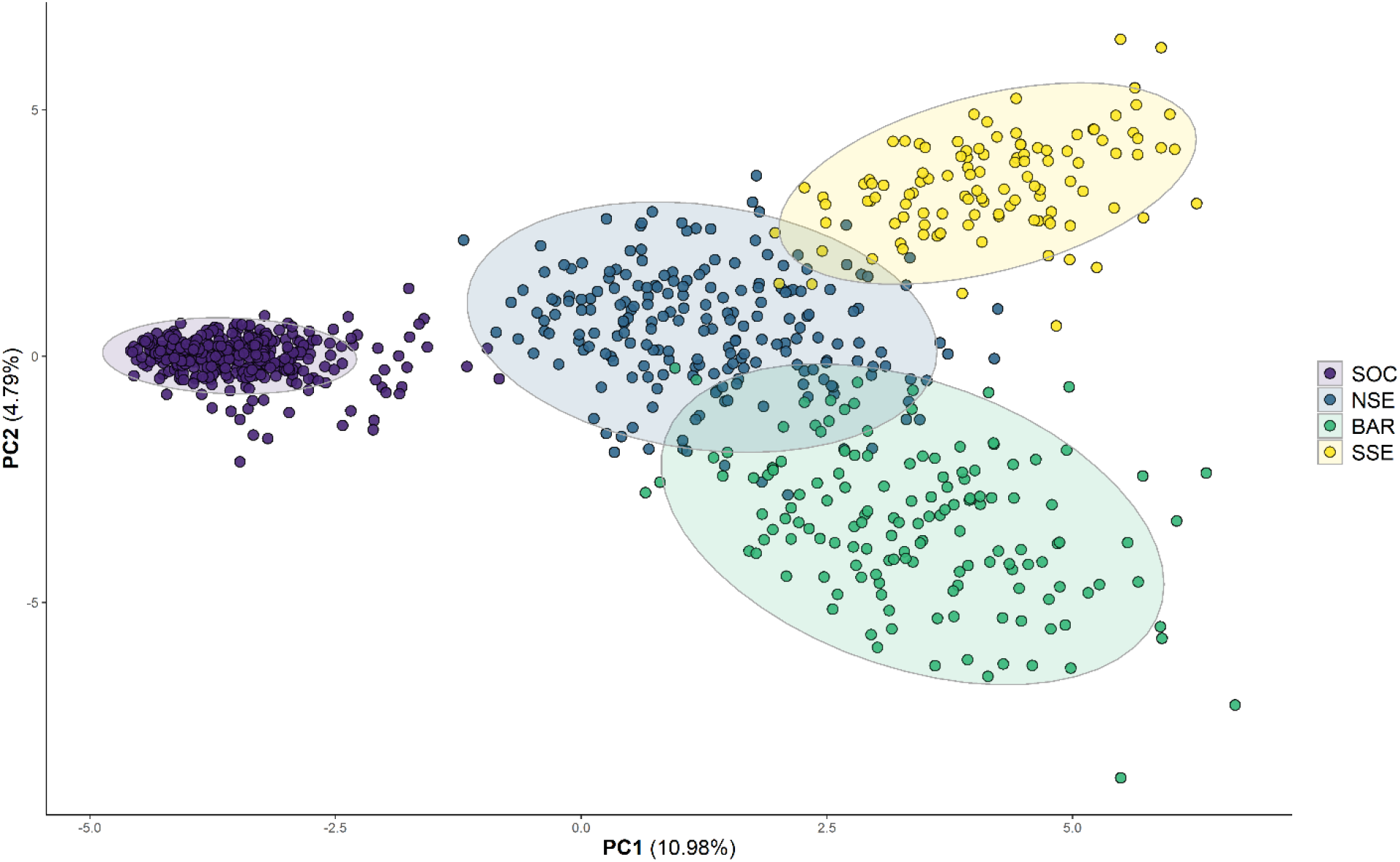
Principal component analysis (PCA) of mountain goat samples in Alaska (n = 816), 2006-2020. The STRUCTURE assignment of each sample is delineated by color. The proportion of variance explained by each axis is shown in parentheses.

**Figure 3.**
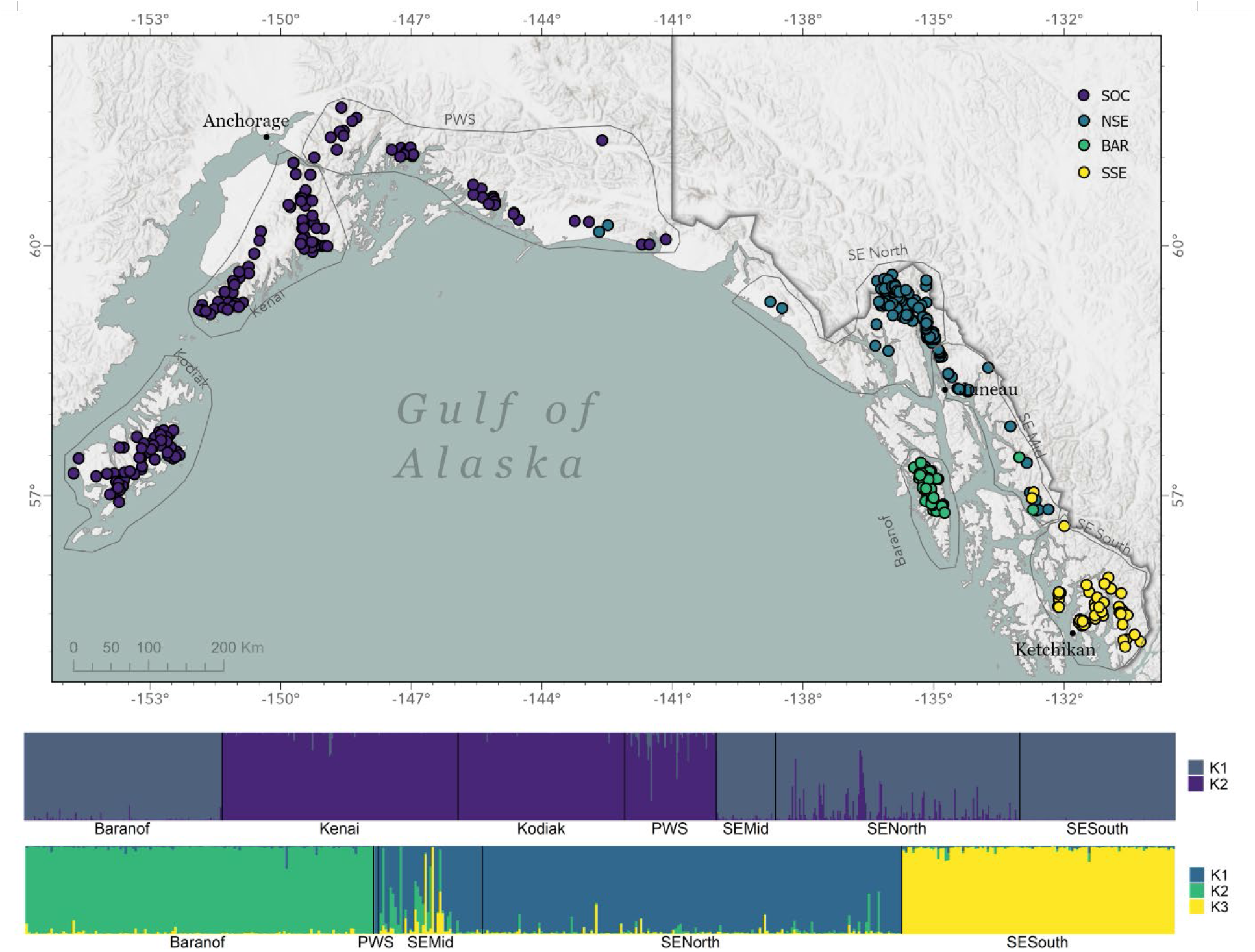
Map of the distribution and location of mountain goat samples (n = 816) collected from 2006-2020 in Alaska, with the assigned STRUCTURE v2.3.4 subpopulation indicated by color. Gray polygons indicate sampling locations. The bar graphs represent the STRUCTURE output for the global run (top) at *K* = 2 and the hierarchical run (bottom) of the Southeast Alaska mountain goats at *K* = 3, ordered by sampling location.

**Figure 4.**
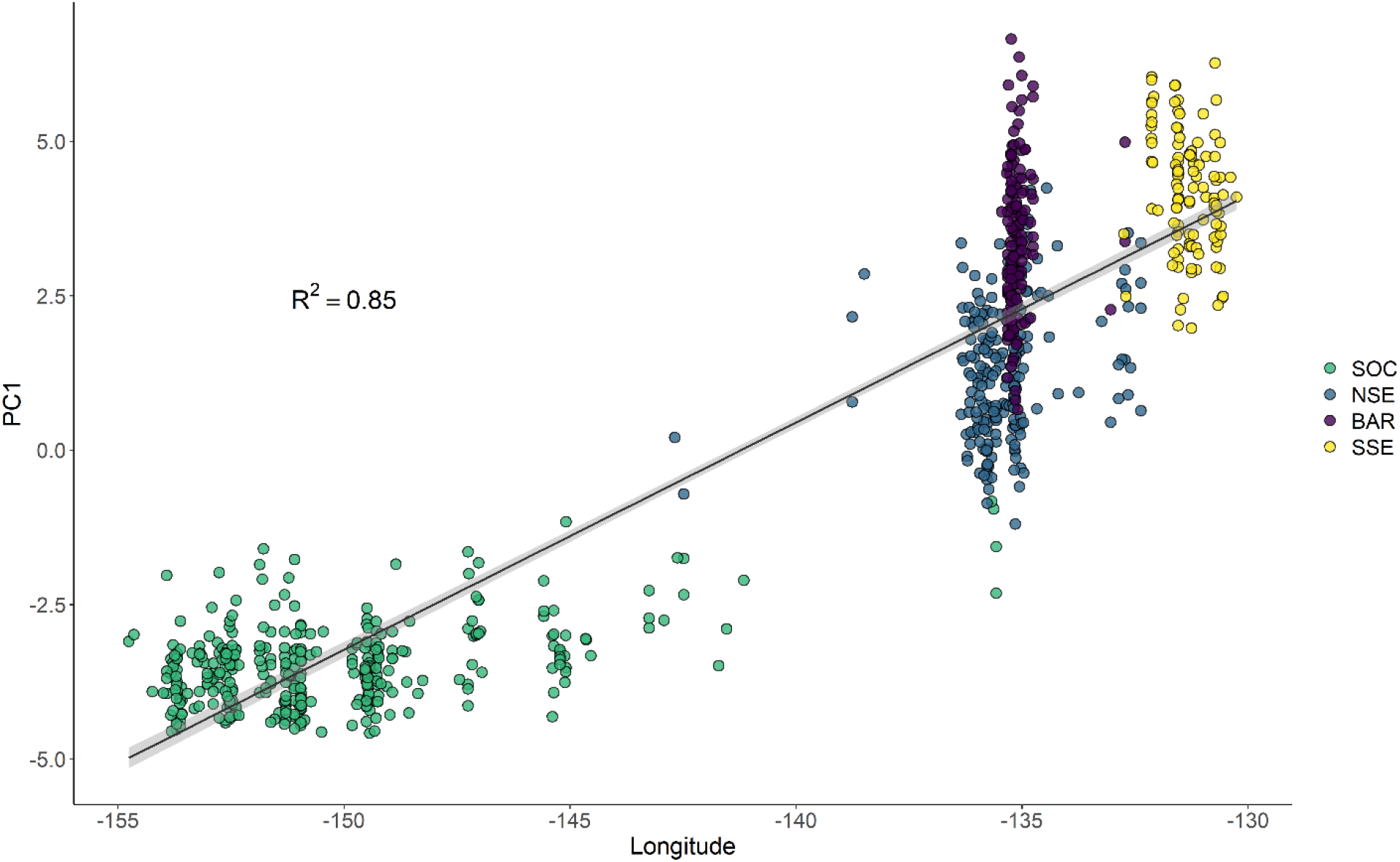
Principle component 1 (PC1) versus longitude for mountain goat allele frequencies (n = 816) in Alaska, 2006-2020. STRUCTURE assignment is indicated by color.

### Diversity Statistics

Nei’s genetic distance analysis showed that subpopulations geographically closest together had the lowest Nei’s *D* and subpopulations farther apart had a higher Nei’s *D*. Nei’s *D* ranged from 0.12 to 0.41 (Table 1). *H_O_* ranged from 0.16 (SOC) to 0.41 (BAR) and *N_e_* ranged from ∼30 to 200. *Fis* ranged from 0.16 (BAR) to 0.29 (SOC) and was positively correlated with latitude (Table 2, Figure S3). In general, SOC had the lowest diversity while the BAR had the highest levels of diversity (Table 2). A Spearman’s rank correlation test showed that Nei’s *D* was highly correlated to *Fst* (ρ = 1, p < 0.01).

**Table 1.**
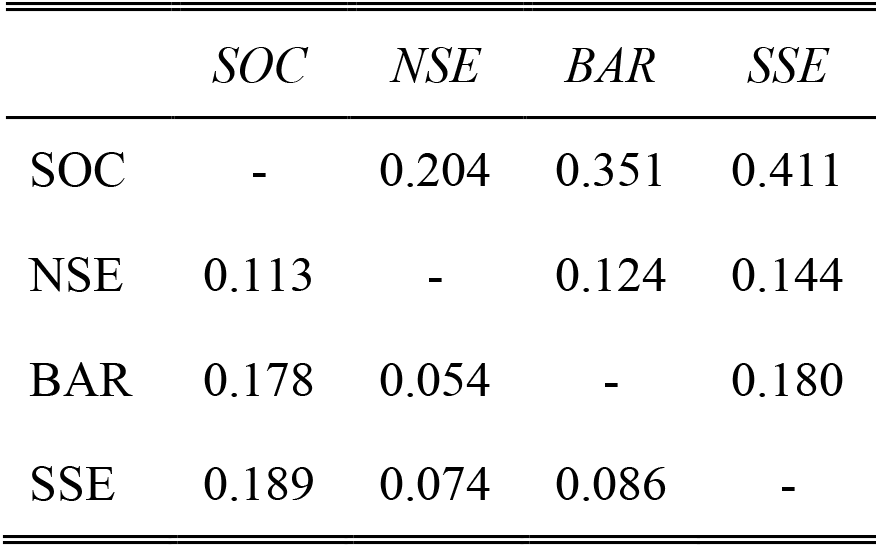
Pairwise *F_ST_* (bottom) and Nei’s *D* (top) for mountain goats in four STRUCTURE defined subpopulations in Alaska (n = 816), 2006-2020. Data were generated using the program SPAGeDi v1.5d and Genalex v6.503.

|  | *SOC* | *NSE* | *BAR* | *SSE* |
| --- | --- | --- | --- | --- |
| SOC | - | 0.204 | 0.351 | 0.411 |
| NSE | 0.113 | - | 0.124 | 0.144 |
| BAR | 0.178 | 0.054 | - | 0.180 |
| SSE | 0.189 | 0.074 | 0.086 | - |

**Table 2.** Diversity statistics for mountain goats in four STRUCTURE defined subpopulations detected in Alaska (n = 1861), 2006-2020. *F_IS_* indicates an inbreeding coefficient and *Ne* indicates the effective population size. Data were generated using Genalex v6.503 and NeEstimator v2.1.

| *Population* | *No. of samples* | $N_e$<br>*95% CI* | *Observed*<br>*heterozygosity* | *Expected*<br>*heterozygosity* | *F<sub>IS</sub>* |
| --- | --- | --- | --- | --- | --- |
| SOC | 352 | 46.6-68.7 | 0.164±0.05 | 0.264±0.05 | 0.287±0.08 |
| NSE | 208 | 77.7-108.8 | 0.362±0.05 | 0.492±0.05 | 0.252±0.05 |
| BAR | 144 | 134.0-360.0 | 0.413±0.05 | 0.483±0.05 | 0.155±0.04 |
| SSE | 112 | 27.2-38.9 | 0.366±0.05 | 0.427±0.06 | 0.139±0.04 |

### Demographic history

We ran the demographic analysis between these Southcentral and Southeast Alaska groupings as per the STRUCTURE *K* = 2 assignment. DIYABC analysis on individuals from each location indicated that Southcentral Alaska mountain goats and Southeast Alaska mountain goats split from each other ∼2,270 generations or ∼13,620 years ago (Figure S4).

### Translocation simulations

The simulated heterozygosity and *F_IS_* for Kodiak and Revilla Islands generally mirrored the observed values for those islands (Table 3). In contrast, the simulated heterozygosity for Baranof Island was significantly smaller than the observed heterozygosity. Allelic richness differed significantly between the simulated and observed individuals on all three islands, though Baranof Island had the largest difference with the observed population having higher allelic richness than the simulated population.

**Table 3.**
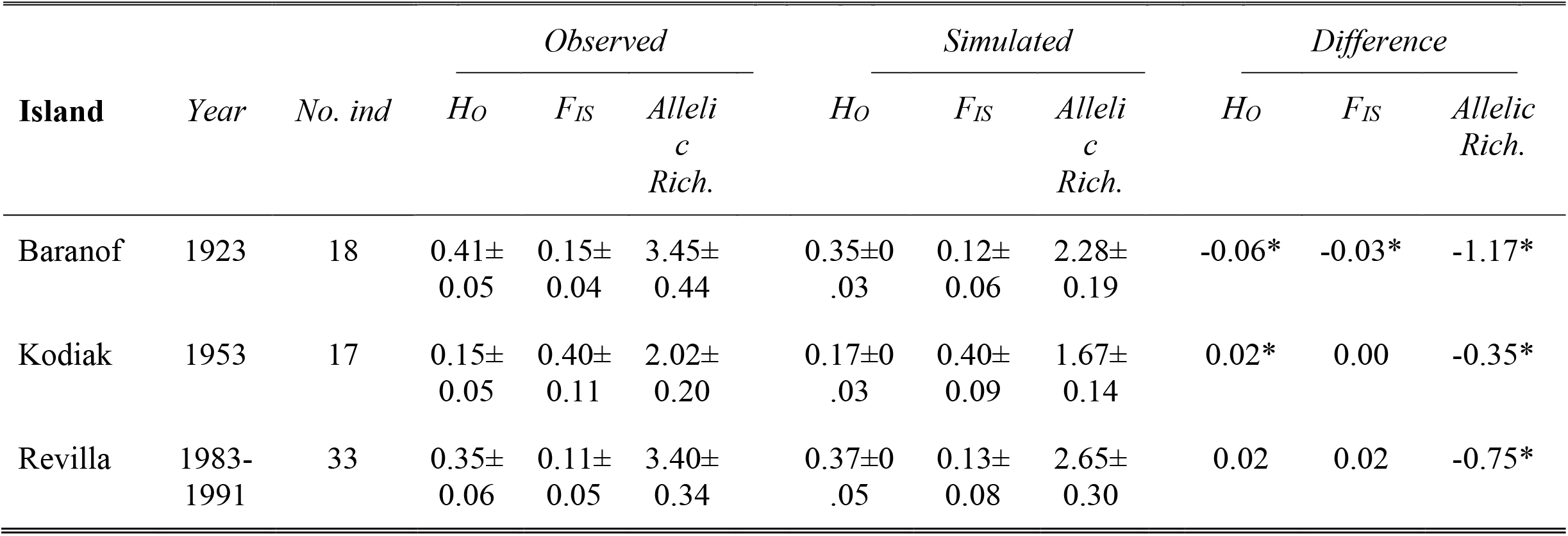
A comparison between the observed and simulated observed heterozygosity (*H_O_*), inbreeding coefficient (*F_IS_*), and allelic richness for mountain goats on three islands in Alaska where translocation events occurred. * indicates significance at p>0.01.

## Discussion

Free-ranging populations are a reflection of the past forces that have acted upon them, both natural and anthropogenic. Genetic variation can record these clues, giving rise to previously unknown refugia (Loehr et al., 2006) and human-assisted translocation events (Chafin et al., 2021; de Jong et al., 2020). Mountain goats are no exception, and our results indicate that their structure is the result of previous glaciations, translocation events, and the current landscape topography. The retreat of the Cordilleran Ice sheet following the end of the Wisconsin glaciation, which began ∼20 thousand calibrated years before present (cal. kyr BP), coincided with the expansion of mountain goats in Alaska. As suitable habitat became ice-free and more widely distributed, mountain goats spread throughout Southcentral and Southeast Alaska to their current distribution. The arrival of European settlers brought an increased demand for hunting and initiated a period where human-assisted translocations occurred. Human intervention, combined with natural factors, further explains the patterns of distribution and structure of mountain goat populations today.

### Demographic patterns and refugial lineages

The Wisconsin glaciation during the late Pleistocene caused large changes in the biodiversity of Alaska, most notably a mass extinction of terrestrial megafauna in North America due to a rapidly changing climate and human impacts (Barnosky et al., 2004; Mann et al., 2015). During the transition period between the Pleistocene and Holocene, temperatures warmed, and plant biodiversity was able to support high numbers of animals, making this a productive time for megafaunal survival and expansion (Guthrie, 2006). Mountain goats were one of the large mammal species in Alaska to survive the mass extinction, and much of the expansion across their range in Alaska occurred during this late Pleistocene and early Holocene period.

Mitochondrial data suggest two main lineages of mountain goats existed prior to the last glaciation, diverging ∼224 cal. kyr BP,in separate refugia (Shafer, Côté, et al. 2011); however, genome-wide data suggest a single refugia during the last glacial maximum followed by northern colonization (Martchenko & Shafer, 2022). Wolf et al. (2020) also suggested a northern expansion following the glacial retreat, and mountain goats in Alaska likely arose from this continued northern expansion. The diversity of Alaskan mountain goats and genetic lineages largely reiterated the geography of their distribution and patterns of IBD (Figure S5). Populations that were farther apart were more genetically distinct from one another compared to neighboring populations. Large mountain ranges and wide fjords influenced population structure with animals in the more naturally fragmented Southeast Alaska region having more distinct populations than Southcentral Alaska. Of the four subpopulations, SOC had generally lower levels of genetic diversity (Table 2). This subpopulation is located geographically far apart from the other subpopulations and is separated by extensive, ice-covered mountains (i.e. the St. Elias Mountains), likely resulting in little to no gene flow between Southeast and Southcentral Alaska.

A variety of other terrestrial mammals in Alaska including moose (*Alces alces*), brown bears (*Ursus arctos*), and gray wolves (*Canis lupus*) have shown a similar pattern of genetic structure where individuals in southeast Alaska are genetically distinct from interior and Southcentral animals, which cluster into one or two genetic populations (Pacheco et al., 2022; Schmidt et al., 2009; Talbot & Shields, 1996). Evidence shows that an ice-free corridor emerged between the Laurentide and Cordilleran Ice sheets around 15-14 cal. kyr BP, allowing for the expansion and divergence of both human and wildlife populations (Heintzman et al., 2016; Loog et al., 2020; Pedersen et al., 2016). Similar timeframes of expansion have been detected globally as was seen with the Northern chamois, a closely related species in the European Alps (Leugger et al., 2022). This timing corresponds with our findings of a northerly mountain goat expansion.

### The impact of translocations

Historically, wildlife translocations have been performed for a variety of reasons including re-establishing locally extinct populations, increasing the genetic diversity of depauperate populations, and for human harvest (Batson et al., 2015; IUCN/SSC, 2013). Regardless of the purpose of the translocation event, consideration of the impacts is important to the future and trajectory of affected populations. Additionally, accounting for past translocations is important in understanding the current structure and conservation status of a population (e.g. Shafer, White, et al., 2011). The three introduced populations of mountain goats provided a unique opportunity to learn about the impact that human intervention can have on the population genetic structure and diversity. Generally, the isolation of mountain goats on islands over the past few decades has not yet created populations that are genetically distinct from their source populations. However, while genetic differentiation is low, it is detected (Table S3), indicating that island mountain goats are starting to drift from their mainland source populations. The rate at which genetic differentiation occurs on isolated island populations depends on a variety of factors including the number, genetic diversity, and age structure of the founding individuals (Le Corre & Kremer, 1998). The exception to this general finding is the mountain goats on Baranof Island, which are genetically distinct from their mainland source populations and show higher levels of diversity than the other two island groups and mainland source (Table 2).

While the Baranof Island translocation occurred before Kodiak and Revilla translocations, this translates to only 5-10 more mountain goat generations. Nevertheless, this difference in time does not explain Baranof Island’s unique level of diversity and differentiation (Tables 1 and 2). Additionally, the simulation analysis, which accounts for the difference in timing, lends evidence that supports Baranof Island having animals present prior to the 1923 translocation event, supporting the existence of an endemic refugial population of mountain goats, as previously reported in Shafer, White et al. (2011) or at least suggests that a single translocation event cannot explain the current patterns. Specifically, when genetic diversity was compared between the simulated and observed populations on the three islands, after parameters were calibrated, the observed genetic diversity was higher than expected with the simulation (Table 3). Translocation events act as a bottleneck and lead to founder effects where a sudden decrease in the number of individuals results in a loss of genetic diversity (Cardoso et al., 2009; Mock et al., 2004). Over time, genetic diversity can be increased through mutation or gene flow, but this process is generally slow, especially on islands where natural gene flow from the mainland is unlikely (Frankham, 1997). If an endemic population of mountain goats already existed on Baranof Island, translocation of animals from an independent source population would facilitate rapid gene flow between the two groups and lead to the observed high rate of genetic diversity of mountain goats on the island.

Glacial refugia have been documented along the North Pacific coast – including areas such as Vancouver Island, that harbored mountain goats (Nagorsen & Keddie, 2000). However, geological evidence of a glacial refugia on Baranof Island documented by Carrara et al. (2007) has recently been questioned (Walcott et al., 2022). Our analyses do not necessarily document evidence of glacial refugia but rather that an endemic population existed prior to the 1923 translocation event. While a glacial refugia represents the most parsimonious explanation for an endemic mountain goat population, the possibility of translocations conducted by indigenous inhabitants of the North Pacific Coast also exists. For example, “gift diplomacy” involving movement of wildlife has been documented among indigenous inhabitants of North America in prehistoric times (Sugiyama et al., 2022), and has been hypothesized to explain the origin of mountain goats on Pitt Island, BC (Menzies 2020, pers. comm). Ultimately, considering both the natural and anthropogenic factors affecting a population is valuable in understanding current population structure and the species history.

## Supporting information

Supplemental Information_Youngetal

## Acknowledgements

We thank J. Breen, E. Wootton, R. Verasztó for their help with DNA extractions and genotyping. Thank you to the helicopter and fixed-wing pilots and ADFG staff who assisted with mountain goat captures and sample collection from harvested animals. This research was funded by the CFI-JELF (36905; A.B.A.S.), Compute Canada Resources for Research Groups (GME-665-01; A.B.A.S.), NSERC Discovery Grant (A.B.A.S.: RGPIN-2017-03934), Ontario Early Researcher Award (A.B.A.S.: ER18-14-209), and the Rocky Mountain Goat Alliance (K.B.Y). This work was conducted in collaboration between Trent University and Alaska Department of Fish & Game.

## Data availability statement

Data used in this analysis have been made publicly available in Dryad.

## Author contributions

KY and AS conceptualized the project. AS and KY obtained funding. KY, KW, and AS collected data and designed methodology. KY analyzed the data. All authors contributed to writing the manuscript.

## Biosketch

K. Young is a wildlife biologist for the Alaska Dept. of Fish & Game. She recently finished her master’s degree studying the population genetics and the effects of climate change and glacial history on mountain goats in Alaska. She is interested in understanding the past and present forces that impact distributions of wildlife species.

K. White is a wildlife research biologist affiliated with the University of Alaska Southeast and University of Victoria, and formerly with the Alaska Department of Fish and Game. He is broadly interested in the population, spatial ecology and conservation of mammals in mountain and northern ecosystems.

A. Shafer is a professor in the Environmental & Life Sciences department of Trent University. He is interested in using genomic tools to study organisms in their natural environment and learning about population processes and histories in hopes of informing conservation and management initiatives.

