## Supplemental Information_Youngetal for "Unravelling the complex biogeographic and anthropogenic history of Alaska’s mountain goats"

### SUPPLEMENTAL TABLES

Table S1. The date, number of individuals, and location of three successful translocations of mountain goats onto islands in Alaska. NR indicates that the number of males and females was not reported for that event.

| *Destination* | *Year* | *Source Pop* | *Males* | *Females* | *Total* | *Reference* |
|---|---|---|---|---|---|---|
| Baranof Island | 1923 | Tracy Arm | NR | NR | 18 | Paul 2009 |
| Kodiak Island | 1952 | Seward | 5 | 0 | 5 | Paul 2009 |
| Kodiak Island | 1953 | Seward | 1 | 9 | 10 | Paul 2009 |
| Kodiak Island | 1952 | Eagle River | 1 | 1 | 2 | Paul 2009 |
| Revillagigedo Island | 1983 | Quartz Hills | 3 | 4 | 7 | Paul 2009 |
| Revillagigedo Island | 1983 | Upp. Cleveland Pen. | 2 | 9 | 11 | Paul 2009 |
| Revillagigedo Island | 1991 | Quartz Hills | NR | NR | 15 | Paul 2009 |

Table S2. Population size for mountain goats on three islands in Alaska where translocations occurred beginning the year of the translocation (Year 0) followed by each subsequent year. Population size data was collected by ADFG during aerial surveys [1-3, 5-18], and we filled in missing data by calculating the average of the known years on either side.

| ***Baranof*** | | | ***Kodiak*** | | | ***Revilla*** | | |
|---|---|---|---|---|---|---|---|---|
| *Year* | *Pop Size* | *# Females* | *Year* | *Pop Size* | *# Females* | *Year* | *Pop Size* | *# Females* |
| 0 | 18 | 8 | 0 | 17 | 10 | 0 | 33 | 17 |
| 1 | 19 | 9 | 1 | 18 | 10 | 1 | 50 | 25 |
| 2 | 20 | 10 | 2 | 18 | 10 | 2 | 66 | 33 |
| 3 | 21 | 11 | 3 | 19 | 10 | 3 | 83 | 41 |
| 4 | 23 | 12 | 4 | 19 | 10 | 4 | 99 | 50 |
| 5 | 25 | 12 | 5 | 20 | 10 | 5 | 119 | 60 |
| 6 | 27 | 13 | 6 | 20 | 10 | 6 | 140 | 70 |
| 7 | 29 | 14 | 7 | 21 | 11 | 7 | 160 | 80 |
| 8 | 30 | 15 | 8 | 21 | 11 | 8 | 166 | 83 |
| 9 | 32 | 16 | 9 | 22 | 11 | 9 | 171 | 86 |
| 10 | 34 | 17 | 10 | 24 | 12 | 10 | 171 | 86 |
| 11 | 36 | 18 | 11 | 26 | 13 | 11 | 172 | 86 |
| 12 | 37 | 19 | 12 | 35 | 18 | 12 | 170 | 85 |
| 13 | 39 | 20 | 13 | 54 | 27 | 13 | 169 | 85 |
| 14 | 41 | 21 | 14 | 58 | 29 | 14 | 162 | 81 |
| 15 | 51 | 25 | 15 | 71 | 36 | 15 | 144 | 72 |
| 16 | 60 | 30 | 16 | 76 | 38 | 16 | 157 | 78 |
| 17 | 70 | 35 | 17 | 81 | 41 | 17 | 169 | 85 |
| 18 | 79 | 40 | 18 | 86 | 43 | 18 | 182 | 91 |
| 19 | 89 | 44 | 19 | 91 | 46 | 19 | 194 | 97 |
| 20 | 98 | 49 | 20 | 112 | 56 | 20 | 207 | 103 |
| 21 | 108 | 54 | 21 | 113 | 57 | 21 | 219 | 110 |
| 22 | 117 | 59 | 22 | 114 | 57 | 22 | 232 | 116 |
| 23 | 127 | 63 | 23 | 115 | 58 | 23 | 315 | 158 |
| 24 | 136 | 68 | 24 | 116 | 58 | 24 | 316 | 158 |
| 25 | 146 | 73 | 25 | 99 | 50 | 25 | 317 | 159 |
| 26 | 155 | 78 | 26 | 120 | 60 | 26 | 360 | 180 |
| 27 | 165 | 83 | 27 | 149 | 75 | 27 | 404 | 202 |
| 28 | 170 | 85 | 28 | 155 | 78 | 28 | 447 | 223 |
| 29 | 175 | 88 | 29 | 175 | 88 | 29 | 490 | 245 |
| 30 | 180 | 90 | 30 | 185 | 93 | | | |
| 31 | 186 | 93 | 31 | 203 | 102 | | | |
| 32 | 191 | 95 | 32 | 360 | 180 | | | |
| 33 | 196 | 98 | 33 | 213 | 107 | | | |
| 34 | 201 | 101 | 34 | 254 | 127 | | | |
| 35 | 206 | 103 | 35 | 196 | 98 | | | |
| 36 | 211 | 106 | 36 | 210 | 105 | | | |
| 37 | 217 | 108 | 37 | 230 | 115 | | | |
| 38 | 222 | 111 | 38 | 250 | 125 | | | |
| 39 | 227 | 113 | 39 | 270 | 135 | | | |
| 40 | 232 | 116 | 40 | 300 | 150 | | | |
| 41 | 237 | 119 | 41 | 325 | 163 | | | |
| 42 | 242 | 121 | 42 | 375 | 188 | | | |
| 43 | 247 | 124 | 43 | 477 | 239 | | | |
| 44 | 253 | 126 | 44 | 596 | 298 | | | |
| 45 | 258 | 129 | 45 | 597 | 299 | | | |
| 46 | 263 | 131 | 46 | 900 | 450 | | | |
| 47 | 268 | 134 | 47 | 1100 | 550 | | | |

| *Baranof* | | | *Kodiak* | | | *Revilla* | | |
|---|---|---|---|---|---|---|---|---|
| *Year* | *Pop Size* | *# Females* | *Year* | *Pop Size* | *# Females* | *Year* | *Pop Size* | *# Females* |
| 48 | 263 | 132 | 48 | 1300 | 650 | | | |
| 49 | 258 | 129 | 49 | 1400 | 700 | | | |
| 50 | 253 | 127 | 50 | 1460 | 730 | | | |
| 51 | 277 | 139 | 51 | 1560 | 780 | | | |
| 52 | 301 | 151 | 52 | 1900 | 950 | | | |
| 53 | 325 | 163 | 53 | 1780 | 890 | | | |
| 54 | 349 | 175 | 54 | 1910 | 955 | | | |
| 55 | 373 | 187 | 55 | 2145 | 1073 | | | |
| 56 | 397 | 199 | 56 | 2371 | 1186 | | | |
| 57 | 473 | 237 | 57 | 2320 | 1160 | | | |
| 58 | 490 | 245 | 58 | 2426 | 1213 | | | |
| 59 | 506 | 253 | 59 | 2390 | 1195 | | | |
| 60 | 515 | 258 | 60 | 2588 | 1294 | | | |
| 61 | 525 | 262 | 61 | 2732 | 1366 | | | |
| 62 | 534 | 267 | 62 | 2732 | 1366 | | | |
| 63 | 530 | 265 | 63 | 3000 | 1500 | | | |
| 64 | 527 | 263 | 64 | 3500 | 1750 | | | |
| 65 | 523 | 262 | | | | | | |
| 66 | 682 | 341 | | | | | | |
| 67 | 841 | 421 | | | | | | |
| 68 | 1000 | 500 | | | | | | |
| 69 | 1050 | 525 | | | | | | |
| 70 | 1100 | 550 | | | | | | |
| 71 | 1150 | 575 | | | | | | |
| 72 | 1200 | 600 | | | | | | |
| 73 | 1250 | 625 | | | | | | |
| 74 | 1300 | 650 | | | | | | |
| 75 | 1350 | 675 | | | | | | |
| 76 | 1350 | 675 | | | | | | |
| 77 | 1350 | 675 | | | | | | |
| 78 | 1350 | 675 | | | | | | |
| 79 | 1350 | 675 | | | | | | |
| 80 | 1440 | 720 | | | | | | |
| 81 | 1788 | 894 | | | | | | |
| 82 | 1755 | 878 | | | | | | |
| 83 | 1722 | 861 | | | | | | |
| 84 | 1689 | 845 | | | | | | |
| 85 | 1656 | 828 | | | | | | |
| 86 | 1623 | 812 | | | | | | |
| 87 | 1590 | 795 | | | | | | |
| 88 | 1557 | 779 | | | | | | |
| 89 | 1524 | 762 | | | | | | |
| 90 | 1491 | 746 | | | | | | |
| 91 | 1458 | 729 | | | | | | |
| 92 | 1431 | 716 | | | | | | |
| 93 | 1324 | 662 | | | | | | |
| 94 | 1521 | 761 | | | | | | |
| 95 | 1717 | 859 | | | | | | |

Table S2. Pairwise $F_{ST}$ (bottom) and Nei's *D* (top) for translocated mountain goat groups from their source population and current island populations. Data were generated using the program SPAGeDi v1.5d. and Genalex v6.503. Bold numbers represent the comparison of the translocated island to its source population.

| | *Baranof* | *BaranofSource* | *Kodiak* | *KodiakSource* | *Revilla* | *RevillaSource* |
|---|---|---|---|---|---|---|
| *Baranof* | - | **0.104** | 0.366 | 0.344 | 0.187 | 0.182 |
| *BaranofSource* | **0.056** | - | 0.240 | 0.232 | 0.146 | 0.124 |
| *Kodiak* | 0.206 | 0.150 | - | **0.026** | 0.429 | 0.412 |
| *KodiakSource* | 0.185 | 0.139 | **0.041** | - | 0.406 | 0.394 |
| *Revilla* | 0.093 | 0.087 | 0.219 | 0.203 | - | **0.010** |
| *RevillaSource* | 0.088 | 0.074 | 0.214 | 0.198 | **0.013** | - |

### SUPPLEMENTAL FIGURES

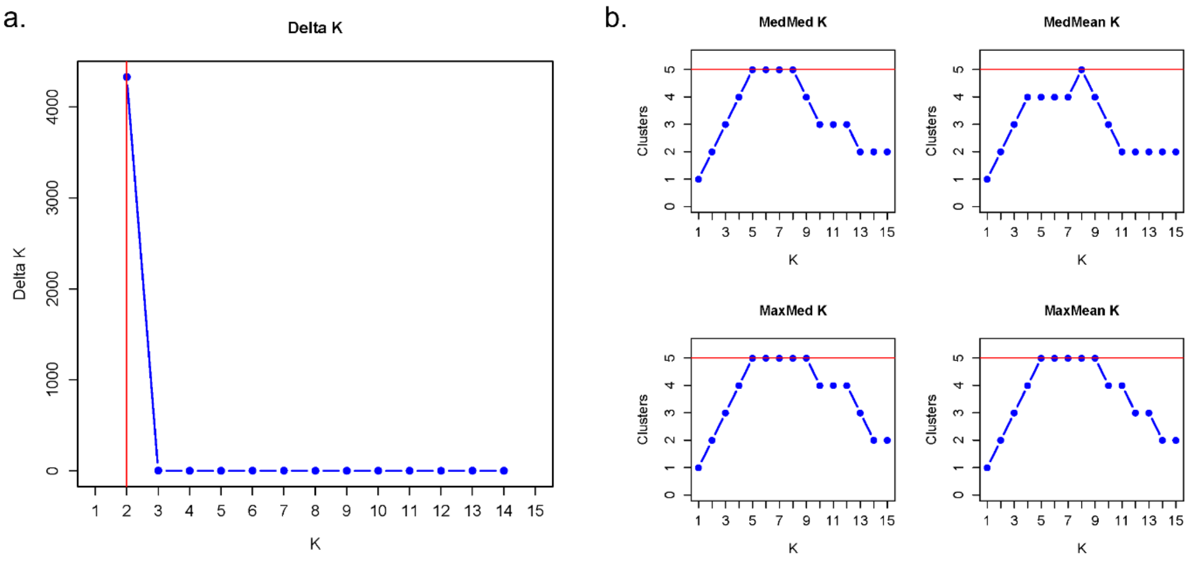

Figure S1. Results from the a) Evanno [4] and b) Puechmaille [19] methods of determining the most likely number of subpopulations of mountain goats in Alaska, 2006-2020. Red lines indicate the most likely number of groupings for each method.

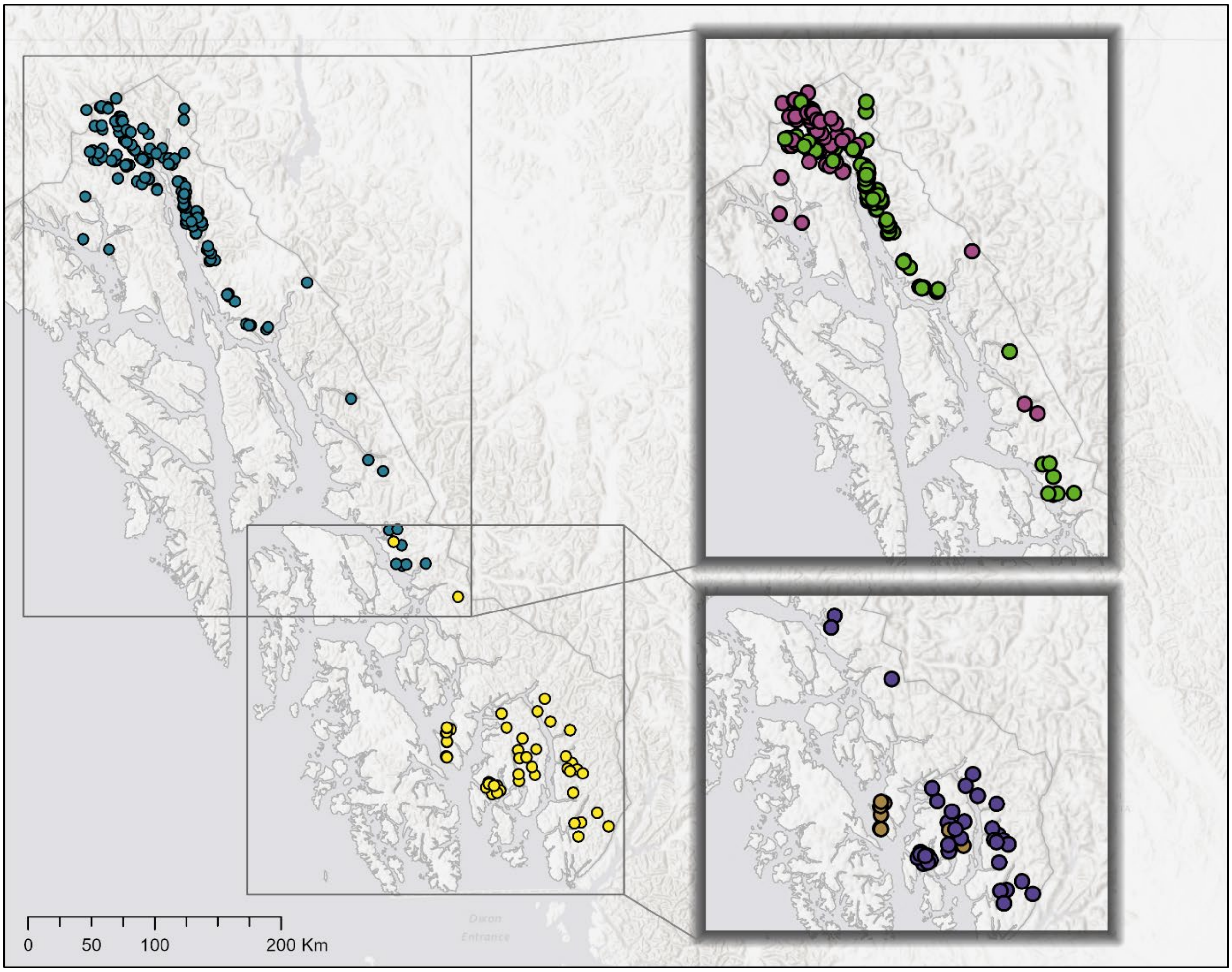

Figure S2. Additional hierarchical structure detected in subpopulations NSE and SSE determined using STRUCTURE v2.3.4 for mountain goats in Southeast Alaska, 2006-2020. Genetically distinct groups are indicated by color.

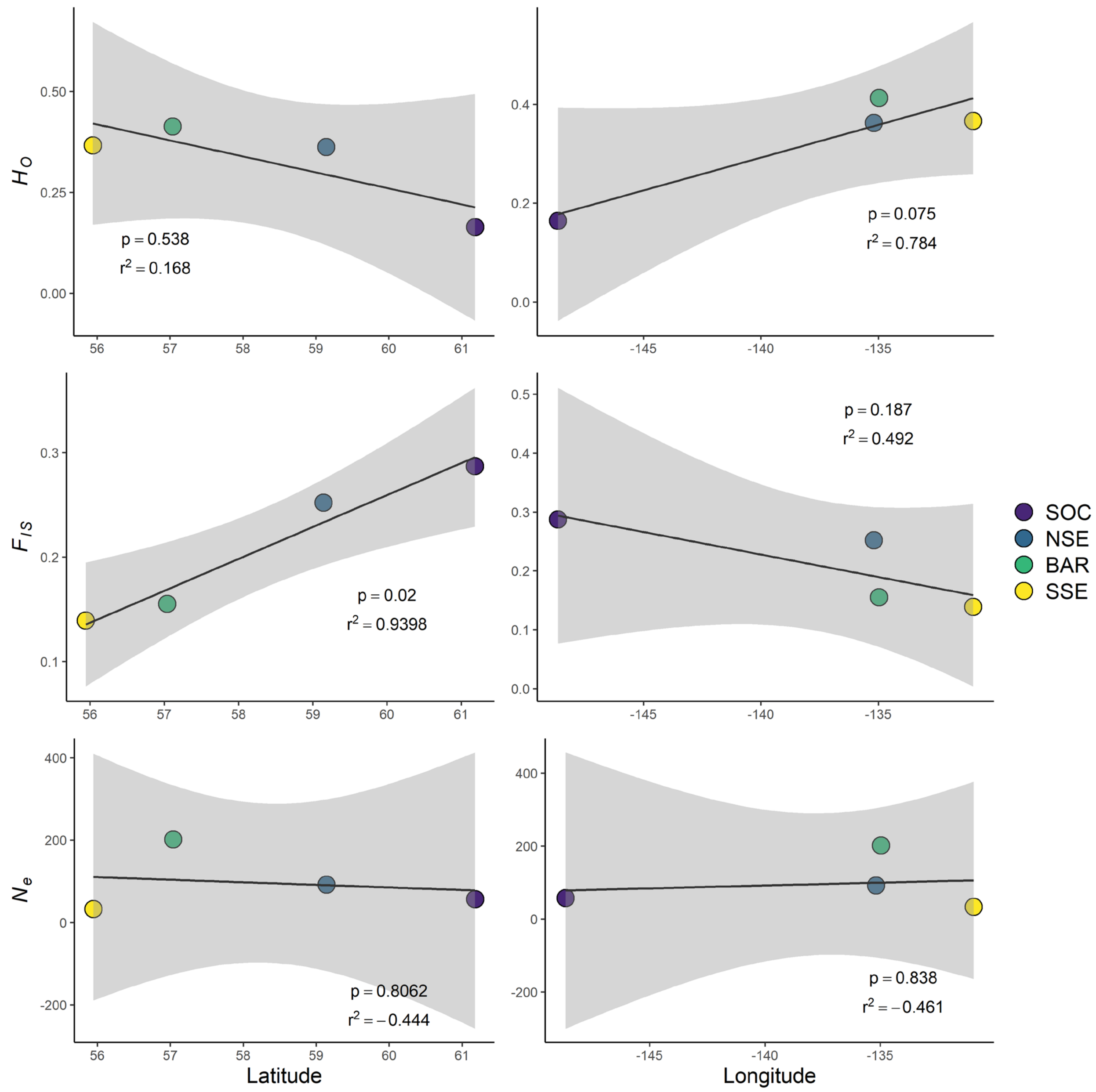

Figure S3. Latitude and longitude plotted against $F_{IS}$, $H_O$, and $N_e$ for mountain goat subpopulations in Alaska (n = 816), 2006-2020. STRUCTURE assigned subpopulations are indicated by color.

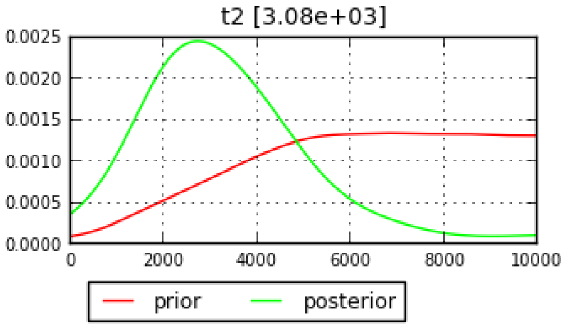

Figure S4. The posterior distribution of split time of mountain goats from southeast and southcentral Alaska, 2006-2020, using the program DIYABC v2.1.0.

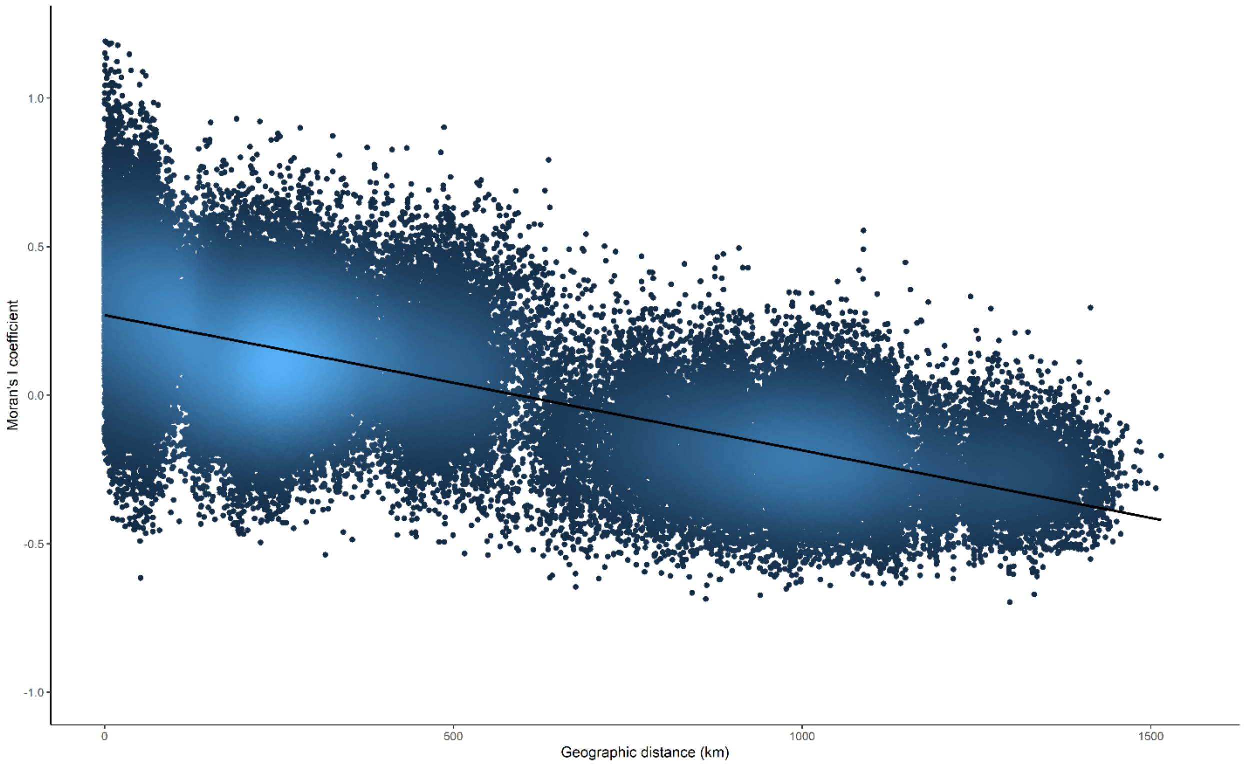

Figure S5. Isolation-by-distance as measured by Moran's I of pairwise genetic relatedness versus the Euclidean geographic distance in kilometers for mountain goats across their range in Alaska (n = 816), 2006-2020.
